# Genome-Wide DNA Methylation Profiles of Neurodevelopmental Disorder Genes in Mouse Placenta and Fetal Brain Following Prenatal Exposure to Polychlorinated Biphenyls

**DOI:** 10.1101/2021.05.27.446011

**Authors:** Benjamin I. Laufer, Kari E. Neier, Anthony E. Valenzuela, Dag H. Yasui, Rebecca J. Schmidt, Pamela J. Lein, Janine M. LaSalle

## Abstract

**Background:** Polychlorinated biphenyls (PCBs) are developmental neurotoxicants implicated as environmental risk factors for neurodevelopmental disorders (NDD), including autism spectrum disorders (ASD).

**Objective:** We examined the effects of prenatal exposure to a human-relevant mixture of PCBs on the DNA methylome of fetal mouse brain and placenta to determine if there was a shared subset of differentially methylated regions (DMRs).

**Methods:** A PCB mixture formulated to model the 12 most abundant congeners detected in the serum of pregnant women from a prospective high-risk ASD cohort was administered to female mice prior to and during pregnancy. Whole-genome bisulfite sequencing (WGBS) was performed to assess genome-wide DNA methylation profiles of placenta and brain on gestational day 18.

**Results:** We found thousands of significant (empirical *p* < 0.05) DMRs distinguishing placentas and brains from PCB-exposed embryos from sex-matched vehicle controls. In both placenta and brain, PCB-associated DMRs were significantly (*p* < 0.005) enriched for functions related to neurodevelopment, cellular adhesion, and cellular signaling, and significantly (Odds Ratio > 2.4, *q* < 0.003) enriched for bivalent chromatin marks. The placenta and brain PCB DMRs overlapped significantly (Z-score = 4.5, *p* = 0.0001) by genomic coordinate and mapped to a shared subset of genes significantly (*q* < 0.05) enriched for Wnt signaling, Slit/Robo signaling, and genes differentially expressed in multiple NDD/ASD models. The placenta and brain DMRs also significantly (*q* < 0.05) overlapped by genomic coordinate with brain samples from humans with Rett syndrome and Dup15q syndrome.

**Discussion:** These results demonstrate that placenta can be used as a surrogate for embryonic brain DNA methylation changes over genes relevant to NDD/ASD in a mouse model of prenatal PCB exposure.

## Introduction

Polychlorinated biphenyls (PCBs) are a class of 209 structurally related congeners that were manufactured in the United States of America beginning in 1929 (Grimm et al. 2015). PCBs were manufactured as a mixture of congeners (e.g. Aroclor) and widely used in electrical equipment, primarily as coolants and insulating fluids for transformers and capacitators, and as stabilizers in a number of commercial products, including paints and caulking. PCB production was banned in 1979 due to concerns about their environmental persistence and human cancer risk (Grimm et al. 2015). Despite the ban, legacy PCBs persist in the environment, and contemporary PCBs not present in the Aroclor mixtures are produced as a by-product of current pigment and dye production used in paints and plastics (Grossman 2013; Hu and Hornbuckle 2010; Kostyniak et al. 2005; Martinez et al. 2012; Sjödin et al. 2014). Contemporary PCBs are detected in both indoor and outdoor air and in human food products (Chen et al. 2017; Hu et al. 2008; Thomas et al. 2012). Due to their persistent and lipophilic nature, legacy PCBs have accumulated in the marine food chains of the great lakes and the Artic, where they place the Indigenous Peoples at elevated risk for exposure (Brown et al. 2018; Hoover et al. 2012; Rawn et al. 2017). Finally, there is documented widespread exposure of humans to PCBs in a number of cities (Hens and Hens 2017).

Prenatal exposure to PCBs can cause developmental neurotoxicity and is considered an environmental risk factor for various neurodevelopmental disorders (NDD), including autism spectrum disorders (ASD) (Klocke et al. 2020; Klocke and Lein 2020; Panesar et al. 2020). Altered DNA CpG methylation, an epigenetic modification, has been associated with PCB exposure and NDDs (Keil and Lein 2016). For example, PCB 95 levels are associated with a gene by environment interaction in a syndromic form of ASD caused by a chromosomal duplication (Dup15q), which is characterized by a global reduction in DNA methylation levels and enrichment for differential methylation at neurodevelopmental genes (Dunaway et al. 2016; Mitchell et al. 2012).

Differential placental DNA methylation has been separately associated with both PCB exposure and NDDs. As the fetal-maternal interface, the placenta is the organ responsible for removing toxicants; however, PCBs are capable of crossing the placental barrier and can also directly impact the placenta (Correia Carreira et al. 2011; Gingrich et al. 2020). In humans, term placenta is accessible at birth and is therefore a potentially rich source of biomarkers for prenatal exposures such as PCBs. PCB exposure was previously shown to be associated with differential methylation at select CpG sites in human placenta in an array-based approach (Ouidir et al. 2020). A low-pass whole genome bisulfite sequencing (WGBS) approach analyzing human placenta samples from a prospective high-risk ASD cohort demonstrated that DNA methylation profiles distinguished ASD from control placenta and the top differentially methylated region (DMR) mapped to CYP2E1 (Zhu et al. 2019). Notably, CYP2E1 plays a key role in the metabolism of PCBs (Chen et al. 2018; Hu et al. 2020; Liu et al. 2017; Uwimana et al. 2019). Furthermore, in the MARBLES cohort, PCBs were detected in the serum of the pregnant mothers at levels that were experimentally shown to impact neurodevelopmental processes in model systems (Sethi et al. 2017a, 2019).

The objective of the research presented in this manuscript was to examine the effect of prenatal PCB exposure on placental and fetal brain DNA methylation profiles from the same mice in a human-relevant exposure model and to determine whether placental DNA methylation patterns could serve as predictors of brain DNA methylation.

## Methods

### Mouse Exposure Model

The PCB mixture formulated to mimic the 12 most abundant congeners identified from the serum of pregnant women in the ASD-enriched MARBLES cohort was prepared as previously described (Sethi et al. 2019). The PCB mixture consisted of the following non-dioxin like congeners in differing proportions: PCB 28 (48.2%), PCB 11 (24.3%), PCB 118 (4.9%), PCB 101 (4.5%), PCB 52 (4.5%), PCB 153 (3.1%), PCB 180 (2.8%), PCB 149 (2.1%), PCB 138 (1.7%), PCB 84 (1.5%), PCB 135 (1.3%) and PCB 95 (1.2%). C57BL/6J dams (The Jackson Laboratory, 000664) aged 6 to 8 weeks were orally exposed to 1.0 mg/kg/d of the PCB mixture through diet (peanut butter) or vehicle (peanut oil in peanut butter) for at least 2 weeks before conception and during pregnancy, as previously described (Keil Stietz et al. 2021). Pregnant dams were euthanized on gestational day 18 and whole brain and placenta were dissected from 44 fetuses and flash frozen. All protocols were approved by the Institutional Animal Care and Use Committee (IACUC) of the University of California, Davis.

### Low-pass WGBS Library Preparation

Nucleic acids were extracted by homogenizing tissues using a TissueLyser II (Qiagen, 85300) followed by the AllPrep DNA/RNA/miRNA Universal Kit (Qiagen, 80224) according to the manufacturer’s instructions. DNA was sonicated to ∼350 bp using a E220 focused-ultrasonicator (Covaris, 500239) and bisulfite converted using the EZ DNA Methylation-Lightning Kit (Zymo Research, D5031) according to the manufacturer’s instructions. Libraries were prepared via the Accel-NGS Methyl-Seq DNA Library Kit (Swift Biosciences, 30096) with the Methyl-Seq Combinatorial Dual Indexing Kit (Swift Biosciences, 38096) according to the manufacturer’s instructions. The library pool was sequenced on all 4 lanes of an NovaSeq 6000 S4 flow cell (Illumina) for 150 bp paired end reads, which yielded ~65 million unique aligned reads (~6X genome cytosine coverage) per a sample.

### Bioinformatic Analyses

The CpG_Me alignment pipeline (v1.4, https://github.com/ben-laufer/CpG_Me), which is based on Trim Galore (v0.6.5), FastQ Screen (v0.14.0), Bismark (v0.22.3), Picard (v2.18.4), and MultiQC (v1.9), was used to trim adapters and methylation bias, screen for contaminating genomes, align to the reference genome (mm10), remove duplicates, calculate coverage and insert size metrics, extract CpG methylation values, generate genome-wide cytosine reports (CpG count matrices), and examine quality control metrics (Ewels et al. 2016; Krueger and Andrews 2011; Langmead and Salzberg 2012; Laufer et al. 2020; Martin 2011; Wingett and Andrews 2018). The raw fastq files and processed CpG count matrices (Bismark genome-wide cytosine reports) are available in the NCBI Gene Expression Omnibus (GEO) repository, through accession number: (available upon publication).

Since PCB exposure and NDDs are known to have substantial sex-specific effects (Keil et al. 2019; Sethi et al. 2017b), the primary analyses were stratified by sex. DMR calling and most downstream analyses and visualizations were performed via DMRichR (v1.6.1, https://github.com/ben-laufer/DMRichR), which utilizes the dmrseq (v1.6.0) and bsseq (v1.22.0) algorithms (Hansen et al. 2012; Korthauer et al. 2018; Laufer et al. 2020). ComplexHeatmap (v2.2.0) was used to create the heatmaps (Gu et al. 2016). GOfuncR (v1.6.1) was used for genomic coordinate based gene ontology (GO) analyses, where DMRs were mapped to genes if they were between 5 Kb upstream to 1 Kb downstream of the gene body, and 1,000 random samplings from the background regions were utilized for the enrichment testing (Grote 2020; Prüfer et al. 2007). HOMER (v4.10) was used to test for the enrichment of transcription factor motifs within the DMRs relative to the background regions using CpG% normalization and the exact sizes of the regions (Heinz et al. 2010). ChIPseeker (v1.22.1) was used to obtain gene region annotations and gene symbol mappings (Yu et al. 2015a). Annotation based enrichment was performed using two sided Fisher exact tests, which was done through LOLA (v1.20.0) for the chromatin state enrichments (Sheffield and Bock 2015), and the odds ratios were converted to fold enrichments for data visualization. enrichR (v3.0) was used for gene symbol based GO, PANTHER pathway, and GEO RNA-seq disease and drug signature enrichment testing (Chen et al. 2013; Jawaid 2021; Kuleshov et al. 2016; Xie et al. 2021). regioneR (v1.22.0) was utilized to perform permutation based genomic coordinate enrichment testing through a randomized region strategy with 10,000 permutations (Gel et al. 2016).

## Results

### Global and Regional CpG Methylation Profiles

To test the hypothesis that prenatal PCB exposure alters DNA methylation profiles in both placenta and embryonic brain, we generated methylomes from matched placenta and brain from PCB-exposed GD18 males (*n* = 11) and females (*n* = 12) and compared to matched vehicle control males (*n* = 10) and females (*n* = 11). The methylomes were profiled by a low-pass WGBS approach that assayed ~20 million CpGs, which is ~ 90% of all CpG sites in the mouse genome. The global methylomes recapitulated known tissue-specific profiles (Schroeder et al. 2015). Specifically, both the female and male placental methylomes (**Figure 1A & 1B**) were hypomethylated when compared to their respective brain methylomes (**Figure 1C & 1D**). In placenta, there was significant global CpG hypomethylation (−1%, *p* ≤ 0.01) in both PCB-exposed females and males when compared to sex-matched controls. Placentas from PCB-exposed females had a global CpG methylation level of 51.9% and control females had a level of 52.8%, while PCB exposed males had a level of 50.5% and control males had a level of 51.5%. In brain, the effects on global methylation were sex-specific, as PCB-exposed males uniquely showed significant global hypermethylation when compared to vehicle male controls (0.1%, *p* = 0.006). Brains from PCB-exposed and control females both had a global CpG methylation level of 76.0%, while brains from PCB-exposed males had a level of 76.2% and control males had a level of 76.1%. In summary, placenta displayed PCB-associated global CpG hypomethylation in both sexes, while only males displayed global CpG hypermethylation in the brain, which had an effect size that was one order of magnitude less than the placental differences.

**Figure 1:**
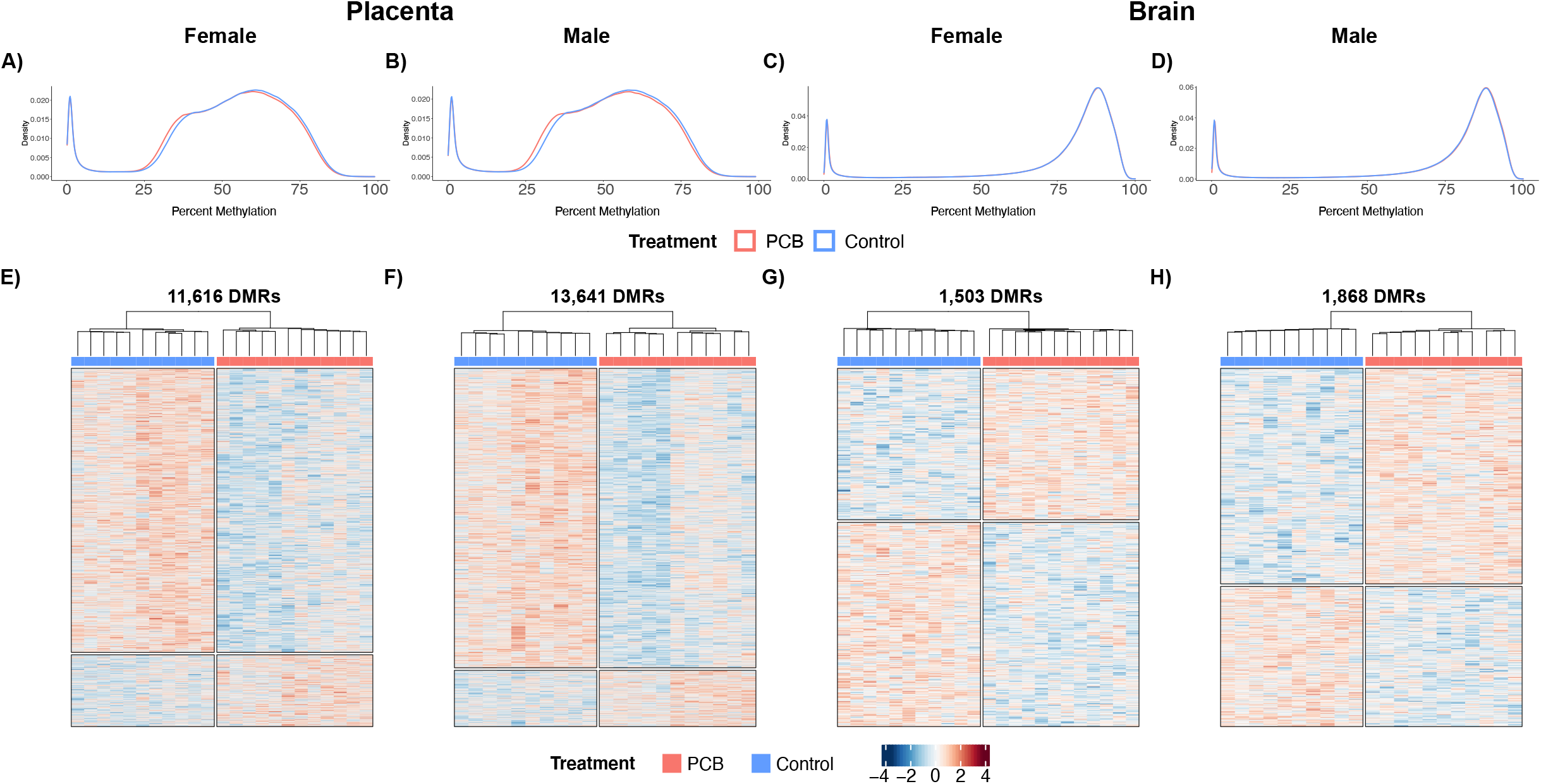
Global and regional DNA methylation profiles of placenta and fetal brain from mice with prenatal PCB exposure. Density plots of smoothed single CpG methylation values from **A)** female placenta, **B)** male placenta, **C)** female brain, and **D)** male brain. Heatmaps of hierarchal clustering of the Z-scores of regional smoothed methylation values for DMRs identified from pairwise contrasts of **E)** female placenta, **F)** male placenta, **G)** female brain, and **H)** male brain.

These differences in global CpG methylation levels were also reflected in finer resolution DMR analyses, which detected significant (empirical *p* < 0.05) genetic locus-specific differences in DNA methylation that distinguished PCB exposed from matched control **(Table 1 & Supplementary Table 1**). In placenta, PCB exposed females displayed a profile of 11,616 DMRs (**Figure 1E**) that were 1,048 bp long and contained 12 CpGs on average. The placenta of PCB exposed males displayed a profile of 13,641 DMRs (**Figure 1F**) that were 1,139 bp long and contained 12 CpGs on average. In brain, PCB exposed females displayed a profile of 1,503 PCB DMRs (**Figure 1G**) that were 568 bp long and contained 11 CpGs on average. Brain from PCB exposed males displayed a profile 1,868 DMRs (**Figure 1H**) that were 608 bp long and contained 10 CpGs on average. In addition to placenta containing approximately an order of magnitude more PCB DMRs than brain, there was a hypomethylation skew in the placental DMRs from both sexes, where 81% of female and 85% of male placental DMRs were hypomethylated. There was a skew towards hypermethylation in only the male brain, where 43% of female and 61% of male brain DMRs were hypermethylated. Finally, the brain DMRs were approximately half the length of the placenta DMRs, on average, despite containing almost the same number of CpGs.

**Table 1:**
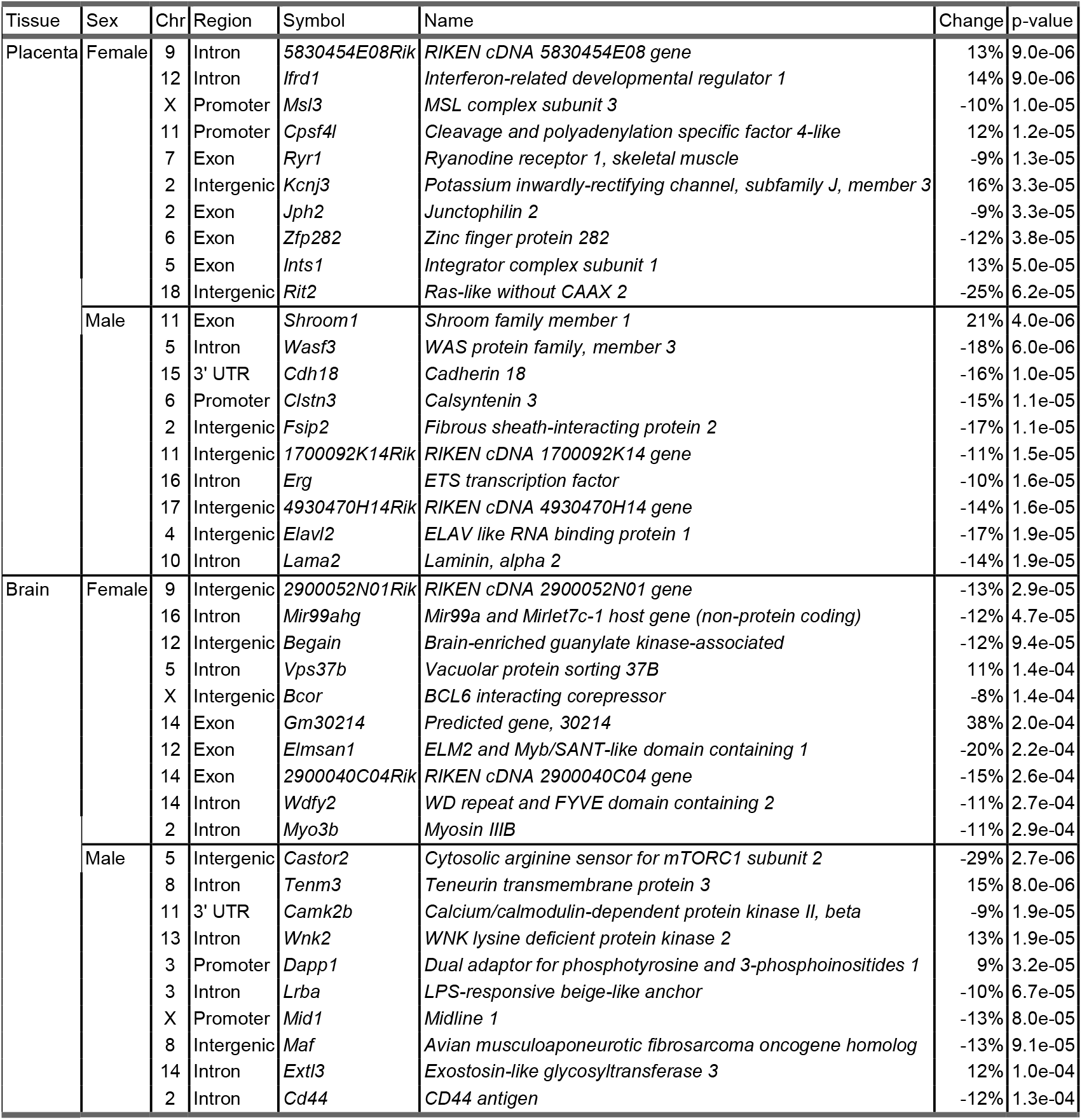
Top 10 DMRs from sex stratified pairwise contrasts of prenatal PCB exposed placenta and fetal brain.

### Functional Enrichments

To test the hypothesis that the prenatal PCB exposure DMRs occur in functional regions of developmentally relevant genes, we performed a series of enrichment testing analyses. We examined the biological relevance of the genes mapping to DMRs through gene ontology (GO) analyses. The top significant (*p* < 0.005) slimmed GO enrichments for female placenta (**Figure 2A**), male placenta (**Figure 2B**), female brain (**Figure 2C**), and male brain (**Figure 2D**) were biological process terms related to neurodevelopment and development, cellular component terms related to the synapse and cell membrane, and molecular function terms related to ion and protein binding as well as cellular signaling (**Supplementary Table 2**). Furthermore, a number of terms passed a more stringent significance (FWER < 0.05) threshold that reduces statistical power, with the placenta showing a stronger enrichment. In female placenta these terms were: protein binding, anatomical structure morphogenesis, nervous system development, binding, cell development, ion binding, synapse, cell projection organization, cell projection. In male placenta the terms were: protein binding, nervous system development, system development, cell adhesion, cell periphery, cell morphogenesis, cell projection organization, localization, ion binding. In female brain the only term was postsynapse and there were none that passed this stringency threshold in male brain.

**Figure 2:**
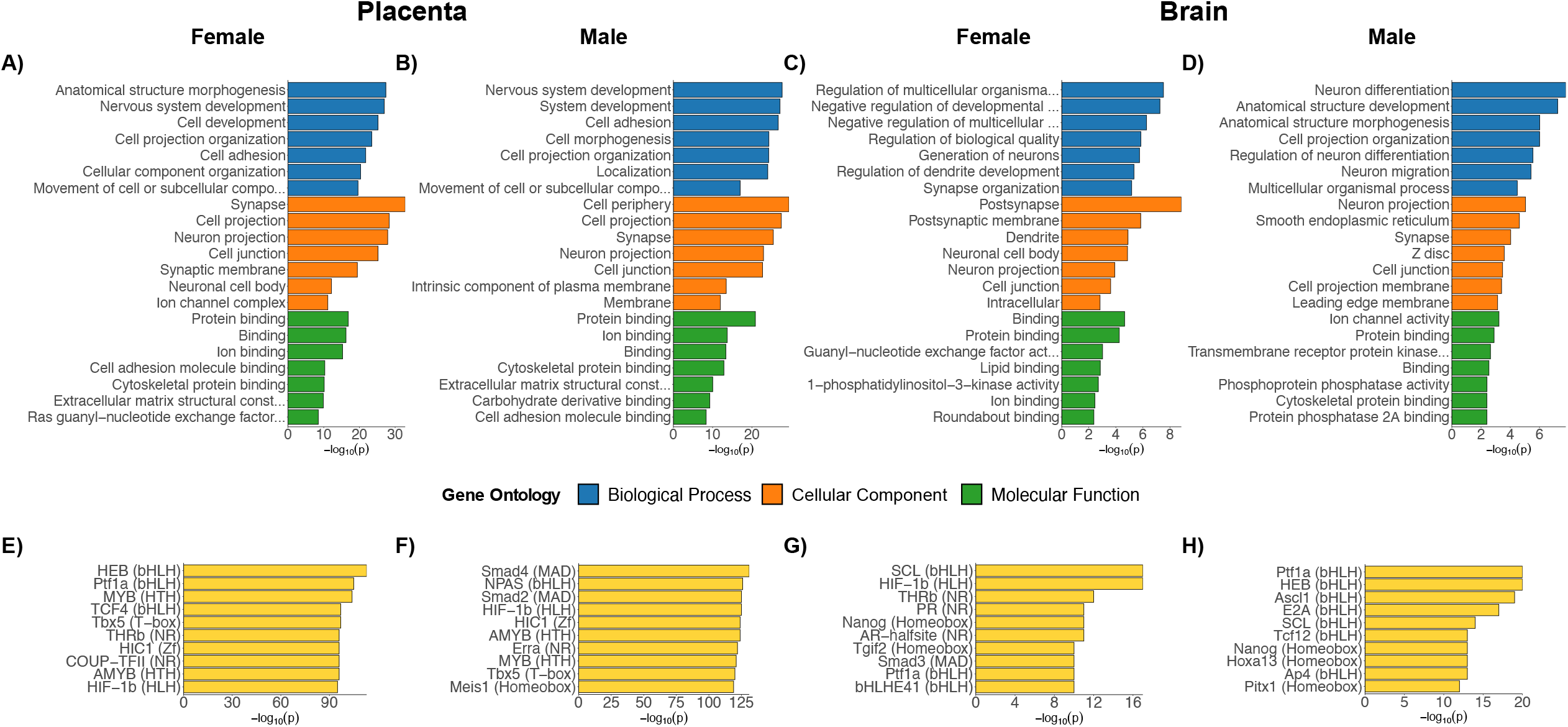
Functional enrichment testing results for prenatal PCB exposure DMRs from placenta and brain. Top slimmed significant (*p* < 0.05) GO enrichment results for DMRs from pairwise contrasts of **A)** female placenta, **B)** male placenta, **C)** female brain, and **D)** male brain. The most significant (*q* < 0.01) transcription factor motif enrichments for pairwise contrasts of **E)** female placenta, **F)** male placenta, **G)** female brain, and **H)** male brain. The motif family is indicated in parenthesis.

To examine the functional gene regulatory relevance of the PCB DMRs, they were tested for enrichment for known transcription factor binding motifs. The GO terms reflecting developmental functions were also reflected in the top significant transcription factor motifs (**Supplementary Table 3**). The most significantly (*q* < 0.01) enriched motifs were: HEB (TCF12) for female placenta (**Figure 2E**), SMAD4 for male placenta (Figure 2F), SCL (TAL1) for female brain (**Figure 2G**), and PTF1A for male brain (**Figure 2H**). Among the top 10 motifs for the different pairwise comparisons, HIF1B (ARNT) and PTF1A were present in 3, while AMYB, HEB (TCF12), HIC1, MYB, NANOG, SCL (TAL1), TBX5, and THRB were present in 2 comparisons.

### Annotation Enrichments

To further test the hypothesis that PCB exposure resulted in methylation differences relevant to gene regulation, the PCB DMRs were tested for enrichment within annotated regions of the genome. The first set of annotation enrichment testing was a tissue agnostic approach to examine CpG and gene region annotations (**Supplementary Table 4**). PCB DMRs were significantly (*q* < 0.05) enriched within CpG islands for both sexes and both tissue sources; however, only the placental PCB DMRs were enriched within CpG shores but depleted within the open sea (**Figure 3A**). PCB DMRs were significantly (*q* < 0.05) enriched within gene bodies but depleted within intergenic regions (**Figure 3B**). The placental PCB DMRs differed from those in brain in that they were also enriched within promoters. Finally, PCB DMRs were tested for enrichment within an 18 chromatin state model of mouse embryonic development, specifically the forebrain tissue (**Supplementary Table 5**) (van der Velde et al. 2021). PCB DMRs from all pairwise contrasts were significantly (*q* < 0.05) enriched within transcription start site (TSS) regions marked by active and bivalent chromatin during at least one developmental timepoint, and bivalent TSS was the top enrichment (Odds Ratio > 2.4, *q* < 0.003) overall (**Figure 3C**).

**Figure 3:**
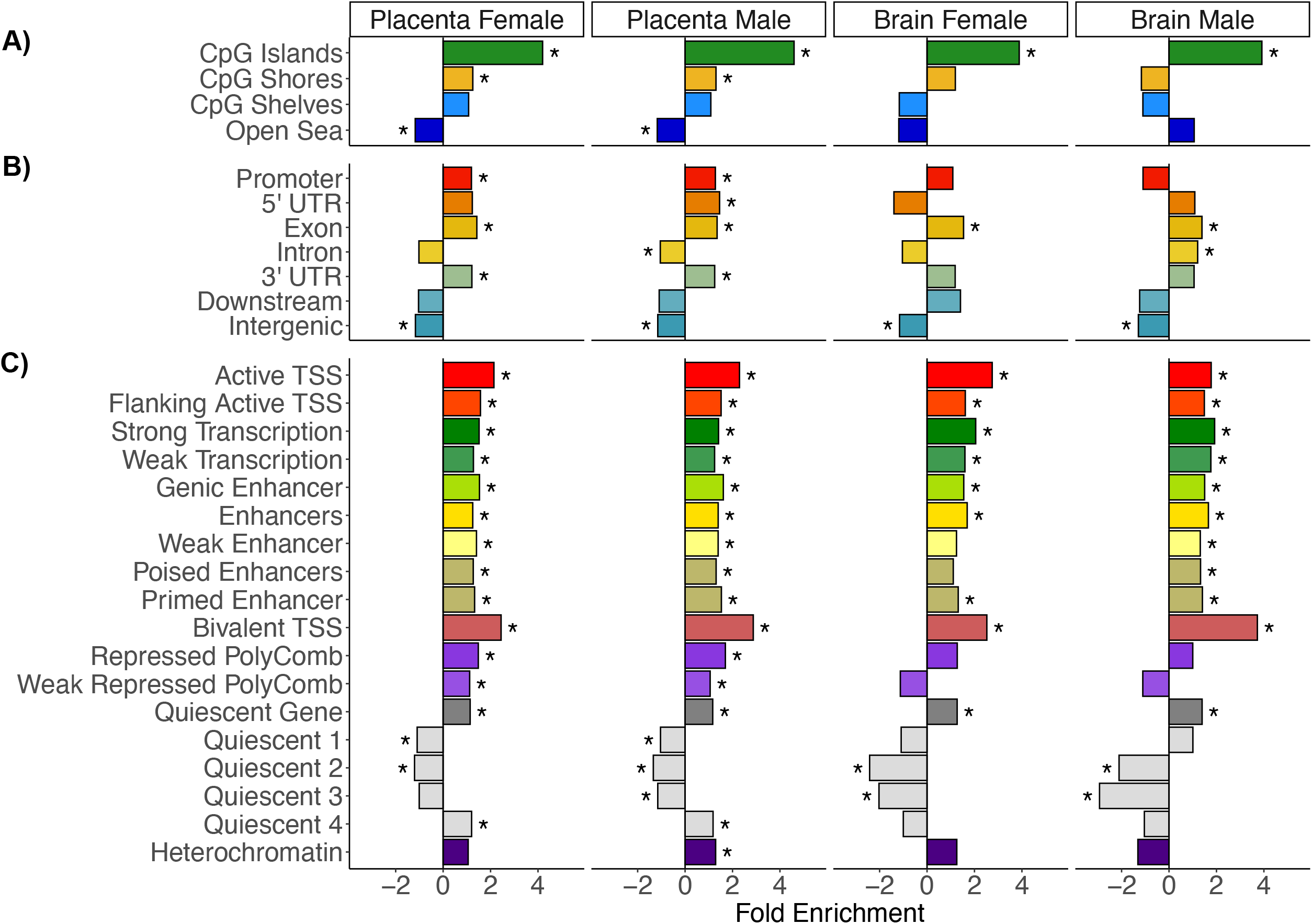
Annotation enrichment testing results for prenatal PCB exposure DMRs from placenta and brain. **A)** CpG annotations, **B)** Gene region annotations, and **C)** Top developmental timepoints for forebrain chromatin states.

### Intersection of Placenta and Brain Methylomes

In order to directly test the significance of overlap between placenta and embryonic brain differential methylation patterns resulting from prenatal PCB exposure, placenta and brain PCB DMRs were examined from both genomic coordinate and gene mapping perspectives. When overlapped by genomic coordinate, the overlapping PCB DMRs mapped to 20 genes in females and 23 in males (**Table 2**). A permutation (*n* = 10,000) based analysis of the genomic coordinate overlap uncovered a significant enrichment for the brain DMRs within the placenta DMRs for females (Z-score = 2.9, *p* = 0.006), males (Z-score = 1.8, *p* = 0.05), and the merging of regions by tissue to produce consensus DMRs (Z-score = 4.5, *p* = 0.0001) (**Figure 4A**). When the DMRs from all pairwise comparisons were mapped to their nearest gene, 210 overlapped by gene symbol (**Figure 4B & Supplementary Table 6**). In order to leverage the statistical power of the sex stratified analyses, a meta p-value analysis was performed on the sex stratified functional enrichment testing results of the gene symbol overlaps between placenta and brain (**Supplementary Table 7**). The top significant (*q* < 0.05) slimmed gene ontology enrichments were primarily related to cell-adhesion, neurodevelopment, metabolism, and cellular signaling (**Figure 4C**). Amongst the top significant (*q* < 0.03) pathways were axon guidance mediated by Slit/Robo, Wnt signaling, and the ionotropic glutamate receptor pathway. In addition to gene functions, this meta-analysis compared gene lists to those from 651 RNA-seq datasets deposited in GEO, which were stratified by direction to provide 1302 gene lists. The top significant (*q* ≤ 6.9×10^−24^) GEO RNA-seq disease and drug signature enrichments were from studies of brain or neuronal responses to stimuli and primarily related to genes repressed by Mecp2 in mouse models of Rett syndrome. There were 86 unique genes shared between both the male and female placenta-brain overlaps and the top GEO datasets (**Supplementary Figure 1**) and 46 of these are from a study of two mouse models of Rett syndrome, specifically the genes repressed by MeCP2 (**Figure 4D**).

**Table 2:**
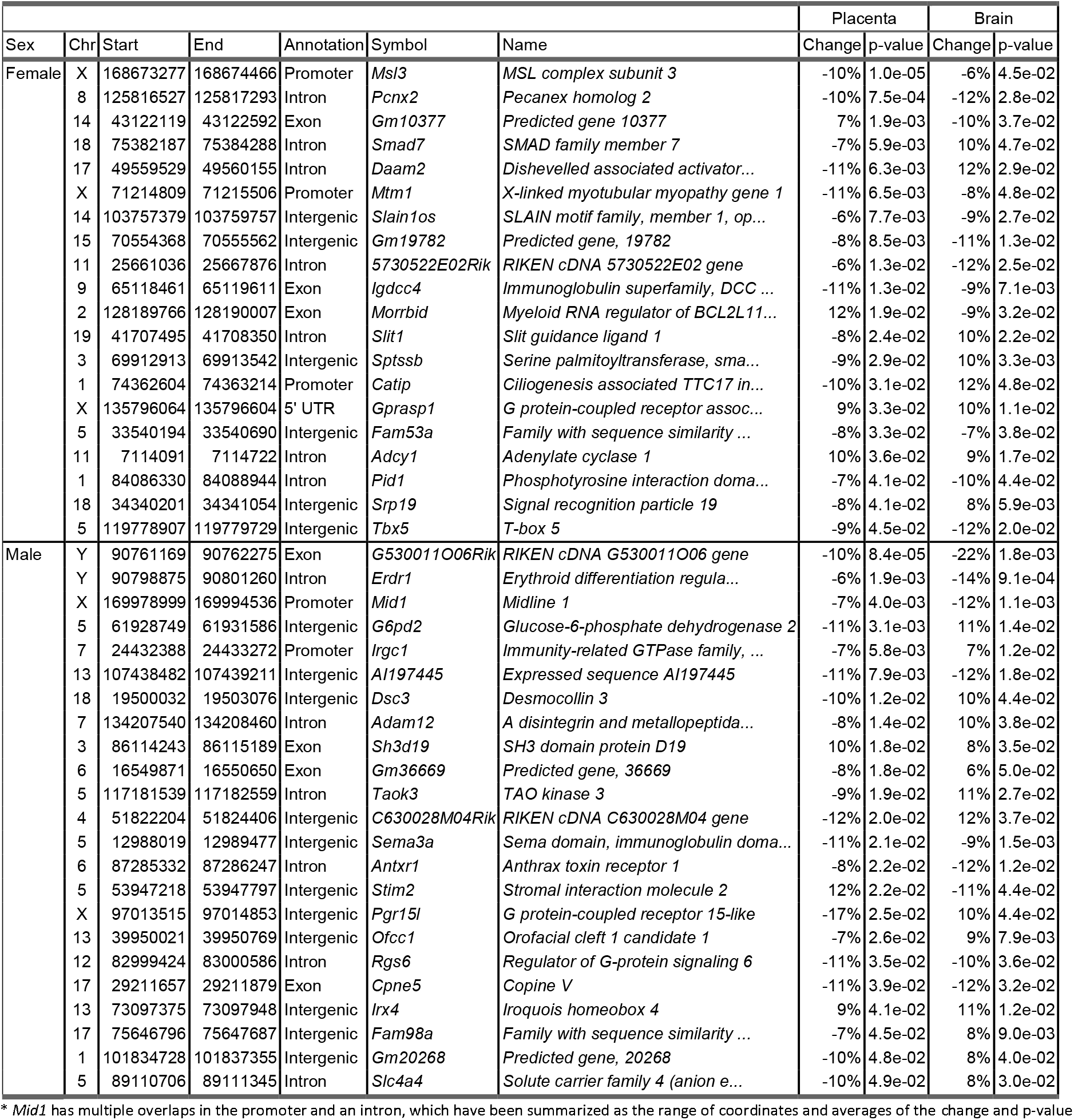
Sex stratified genomic coordinate overlaps for prenatal PCB exposure DMRs from placenta and fetal brain.

**Figure 4:**
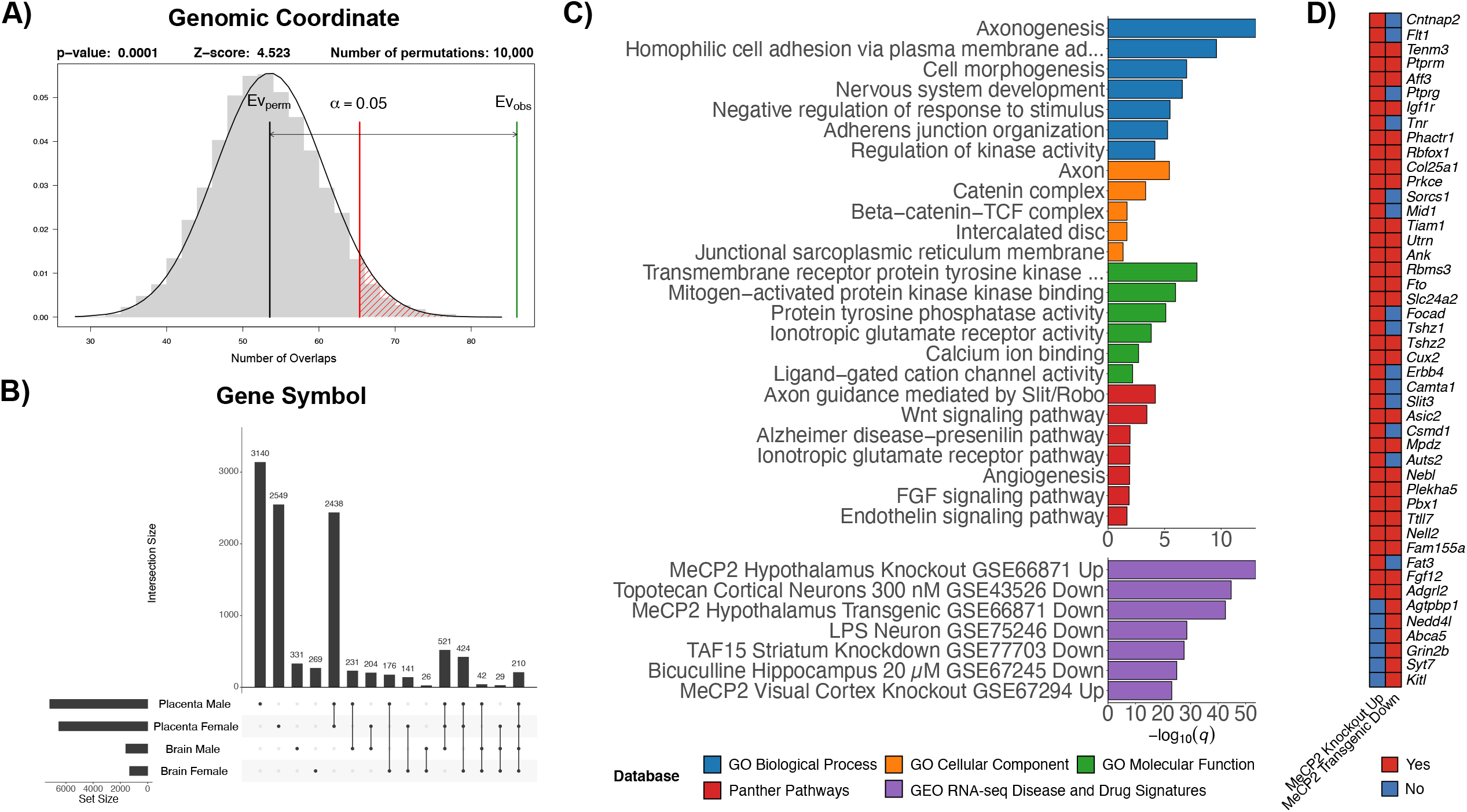
Overlaps between PCB exposure DMRs from placenta and brain. A) Permutation analysis of the genomic coordinate enrichment of the sex combined brain DMRs within the sex combined placenta DMRs. B) UpSet plot of the gene symbol mapping overlaps for all pairwise DMR contrasts. C) Top significant (*q* < 0.05) slimmed GO terms, Panther pathways, and GEO RNA-seq dataset enrichments from a meta p-value analysis of genes symbol mappings from the sex stratified overlaps of placenta and brain. The GEO RNA-seq dataset enrichments are stratified by whether the genes were up- or down-regulated. D) Heatmap of unique DMR gene symbol mappings that are shared between the female and male placenta-brain overlaps and repressed by MeCP2 in the hypothalamus of mouse models of Rett syndrome.

Next, we examined the relevance of the findings to humans by testing the PCB DMRs for enrichment within regions identified by human epigenome-wide association studies (EWAS). First, we tested the hypothesis that that PCB exposure DMRs are enriched within differentially methylated CpG sites identified by the Infinium Methylation EPIC BeadChip in humans with PCB exposure (Curtis et al. 2020; Pittman et al. 2020). Since the human studies analyzed both sexes together, the sex combined tissue consensus DMRs were tested after being lifted over to the human genome. Only the consensus brain DMRs were significantly (Z-score = 6.7, *q* = 0.001) enriched within sites associated with PCB levels in human peripheral blood samples (Curtis et al. 2020). Second, we tested the hypothesis that the PCB exposure DMRs were enriched within DMRs associated with NDDs identified from brain. We utilized two of our previously published brain NDD DMR datasets, after updating them to be processed similarly, including male patients with chromosome 15q11.2−13.3 duplication syndrome (Dup15q syndrome) and high PCB levels as well as female patients with Rett syndrome (Dunaway et al. 2016; Vogel Ciernia et al. 2020). The consensus brain PCB associated DMRs were significantly enriched within the Dup15q syndrome (Z-score = 5.1, *q* = 0.0004) and Rett syndrome brain DMRs (Z-score = 2.4, *q* = 0.02) and the consensus placenta PCB associated DMRs were also significantly enriched within the Dup15q syndrome (Z-score = 5.9, *q* = 0.0004) and Rett syndrome (Z-score = 4.7, *q* = 0.0004) brain DMRs.

## Discussion

This epigenomic study provides multiple novel insights into the impact of prenatal PCB exposure on neurodevelopment by examining both placenta and embryonic brain in a human-relevant exposure model in mice. First, the results demonstrate that prenatal PCB exposure results in sex-specific genome-wide DNA methylation differences in gene regulatory regions in both placenta and fetal brain. Interestingly, the impact of prenatal PCB exposure on the epigenome was stronger in placenta than brain and a subset of DMRs from placenta mapped to genes enriched for neurodevelopmental functions, including synaptic, cell adhesion, and ion binding functions. Second, the differentially methylated genes in both placenta and brain were associated with known NDDs and reflect cellular signaling pathways that have been previously shown to be disrupted in brain cells by developmental PCB exposure. These results suggest that human term placenta, a tissue accessible at birth, could be utilized for identifying epigenetic biomarkers of prenatal PCB exposure in the fetal brain.

PCBs can be divided into two categories based on their mechanisms of toxicity: dioxin-like (DL) and non-dioxin like (NDL). However, there are similarities in that developmental exposure to DL and NDL PCBs has been shown decrease levels of thyroid hormone in serum (Bansal et al. 2005; Gauger et al. 2004; Giera et al. 2011; Zoeller 2007). Thyroid Hormone Receptor Beta (THRB) was among the top 10 transcription factor motifs enriched within placenta and brain PCB DMRs in females, with lower ranked enrichments in males. DL PCBs differ from NDL PCBs in that their primary mechanism involves their binding to the aryl hydrocarbon receptor (AHR), which is then bound by the aryl hydrocarbon receptor nuclear translocator (ARNT), also known as hypoxia-inducible factor 1β (HIF1B), and translocated to the nucleus to activate genes involved in xenobiotic metabolism, such as the such as cytochrome P450s (Calò et al. 2014; Kim et al. 2015; Klinefelter et al. 2018; Matsushita et al. 1993; Seok et al. 2017). HIF1B was one of the top 10 transcription factor motifs enriched in PCB DMRs from female placenta, male placenta, and female brain, with lower ranked enrichment in male brain. Furthermore, a cytochrome P450 (CYP2E1) was one of the top ASD-associated DMRs in placental samples from the MARBLES study (Zhu et al. 2019), which was the reference for the PCB congener mixture used in this study (Sethi et al. 2019). Genes encoding multiple additional members of the cytochrome P450 family were also present in the placental PCB DMRs from our current study. Together, these results implicate known targets of DL PCBs and those shared with NDL PCBs; however, most of the DNA methylation differences observed with PCB exposure were related to the known mechanisms of the NDL PCBs.

Legacy and contemporary NDL PCBs predominately act through calcium-dependent mechanisms, where they increase intracellular calcium levels to alter synaptic connectivity (Klocke et al. 2020; Klocke and Lein 2020). Neurodevelopmental terms related to synaptic connectivity and terms related to calcium ion binding were present in the top terms for the pairwise GO analyses as well as the GO terms for the overlap between placenta and brain. The effects on calcium signaling are driven by legacy NDL PCBs activating signaling proteins on the cell membrane and endoplasmic reticulum, which were also enriched within both pairwise and placenta-brain overlap GO analyses. At the cell membrane, PCBs activate NMDA receptors, a type of ionotropic glutamate receptor, and L-type voltage-gated calcium channels (Inglefield and Shafer 2000; Llansola et al. 2009, 2010). Glutamate receptor functions were enriched within both GO molecular function and PANTHER pathway terms for placenta-brain overlaps. PCBs also act on ryanodine and inositol 1,4,5-tris-phosphate receptors within the endoplasmic reticulum membrane (Inglefield et al. 2001; Pessah et al. 2010). Amongst all the membrane proteins targeted by legacy NDL PCBs, the most responsive are ryanodine receptors, which become sensitized (Klocke et al. 2020; Klocke and Lein 2020; Panesar et al. 2020; Pessah et al. 2010). PCB 136 has been shown to sensitize ryanodine receptors and increase the frequency of spontaneous calcium oscillations in primary cultures of rat hippocampal neurons (Yang et al. 2014). In our study, ryanodine receptor 1 (Ryr1) was the 5^th^ highest ranked PCB DMR in female placenta and each pairwise PCB comparison contained more than one DMR mapping to a ryanodine receptor gene. The endoplasmic reticulum was also one of the top cellular component GO terms in male brain, and was also represented in the placenta and brain overlap cellular component GO terms.

The disruption to calcium signaling via the ryanodine receptor results in the deregulation of downstream developmental pathways. In primary rat hippocampal cultures, PCB 95 exposure has been shown to potentiate ryanodine receptor calcium channels, leading to increased calcium oscillations that activate the calcium/calmodulin-dependent protein kinase type 1 (CaMKI) and result in the CREB promoting transcription of Wnt2 to ultimately stimulate dendritic growth and synaptogenesis (Lesiak et al. 2014; Wayman et al. 2012a, 2012b). The effects of the CaMKI signaling cascade on Wnt signaling appear to be the primary pathway reflected in the DMRs that overlap between placenta and brain. The GO terms and pathways for the genes shared between placenta and brain primarily represent a cascade related to the cadherin pathway, which mediates calcium ion dependent cell adhesion. β-catenin is a subunit of the cadherin complex that functions as an intracellular signal transducer for the Wnt signaling pathway (Steinhart and Angers 2018). The Wnt pathway is also represented in the placenta and brain genomic coordinate overlapped DMRs in females through Daam2 and through Wnk2 in the male brain DMRs (Lee and Deneen 2012; Serysheva et al. 2013). Although some contemporary NDL PCBs are not as well studied given their recent emergence, PCB 11 has been shown to effect dendritic arborization through a CREB-dependent mechanism in primary rat cortical neuron-glia co-cultures (Sethi et al. 2018). Therefore, while there are differences in the mechanisms of some legacy and contemporary NDL PCBs, they converge at cAMP response element-binding protein (CREB) signaling. In our study, CREB signaling is represented by Adenylate cyclase 1 (Adcy1) in the female placenta and brain genomic coordinate overlapped DMRs and by Calcium/calmodulin-dependent protein kinase II, beta (Camk2b) in the top male brain DMRs. Overall, the results demonstrate that disruptions to known PCB mediated signaling cascades are reflected in the brain and placenta methylome.

The DNA methylation profile of prenatal PCB exposure also refines the effects on neurite outgrowth to the Slit/Robo signaling pathway. This pathway consists of the secreted Slit proteins and their receptors, the Roundabout (Robo) proteins. Although initially characterized for their role in axon guidance, members of Slit/Robo signaling are involved in dendritic growth and branching (Whitford et al. 2002). Slit/Robo signaling was the top ranked PANTHER pathway in the analysis of placenta and brain overlaps and roundabout binding was one of the top GO terms for female brain. This signaling pathway is represented in the genomic coordinate overlaps between placenta and brain through Slit1 in females and Mid1 (Midline 1) in males (Liu et al. 2009; Whitford et al. 2002). Interestingly, many of the pathways that are downstream or cross-talk with Slit/Robo signaling in diverse cell types are represented in the prenatal PCB exposure DMRs, which includes Wnt, PI3K/AKT/mTOR, and TGF-β signaling (Blockus and Chédotal 2016). PCB 95’s promotion of dendritic growth involves mTOR signaling in primary rat hippocampal neuron-glia co-cultures (Keil et al. 2018). The PI3K/AKT/mTOR pathway was reflected in the female brain DMRs through the 1−phosphatidylinositol−3−kinase activity molecular function GO term and in male brain through the top DMR, which mapped to Cytosolic arginine sensor for mTORC1 subunit 2 (Castor2). TGF-β signaling is represented in the results through its signal transducers: the Smad proteins. Smad7 overlapped by genomic coordinate in female placenta and brain PCB DMRs, and Smad proteins were among the top transcription factor motif enrichments for male placenta and female brain. Thus, it appears that many of intracellular signaling cascades that are downstream or cross-talk with Slit/Robo signaling are represented in the prenatal PCB exposure DMRs. In addition to a critical role in neurodevelopment, the above signaling cascade also appears to reflect the anti-angiogenic effects of PCBs on placenta (Kalkunte et al. 2017). Daam2, a member of the Wnt signaling pathway, is involved in placental vascularization (Nakaya et al. 2020). The Slit/Robo pathway has also been implicated in placental angiogenesis (Bedell et al. 2005; Chen et al. 2016; Liao et al. 2012). The effect on angiogenesis and vascularization is also apparent in the placenta-brain overlapping DMRs through the angiogenesis and endothelin signaling PANTHER pathway enrichments. Taken together, these findings show that the disrupted signaling pathways have distinct functions in both neurodevelopment and placental development.

Aside from being enriched within known signaling pathways, the genes mapping to PCB DMRs are strongly enriched for transcriptional dysregulation in neurodevelopmental disorders and neuronal drug responses from previously published datasets. The most prominent of the enrichments was for genes repressed by methyl-CpG binding protein 2 (MeCP2) in brain from mouse models of Rett syndrome (Chen et al. 2015; Gabel et al. 2015). One of these studies demonstrated that MeCP2 represses the expression of long genes enriched for ASD risk (Gabel et al. 2015). The NDD signature related to long genes and ASD risk is also apparent through an enrichment for genes down-regulated by topotecan, a topoisomerase inhibitor that reduces expression of many long genes associated with ASD in neurons (King et al. 2013). The disease and drug signature analysis also identified two other treatments that have been previously implicated in PCB exposure. Bicuculline is a GABA receptor agonist (Yu et al. 2015b), which has been shown to phenocopy the effects of PCB-95 on dendritic growth (Wayman et al. 2012a). There was also an immune signature represented through lipopolysaccharide (LPS) challenge in neurons (Srinivasan et al. 2016). This enrichment is further reflected by the top female brain DMR mapping to 2900052N01Rik, which shows high expression in B cells that have been stimulated by LPS (Wang et al. 2021), and through LPS-responsive beige-like anchor (Lrba), which was one of the top DMRs male brain. Additionally, there are a number of immune genes in the top DMRs as well as the placenta and brain overlapped DMRs associated with PCB exposure. Finally, there was a signature related to neurodegenerative diseases characterized by neuritic plaques and neurofibrillary tangles, specifically through a study of the role of TAF15 in amyotrophic lateral sclerosis (ALS) and the Alzheimer disease−presenilin pathway from PANTHER (Kapeli et al. 2016). Ultimately, along with the genomic coordinate based enrichment of the consensus brain PCB associated DMRs within differentially methylated sites from human lymphocytes with measured PCB exposures and the enrichment of the PCB associated DMRs from both placenta and brain within brain from individuals with Rett and Dup15q syndrome, the prenatal PCB exposure DMRs identified in mouse show a profile that is relevant to human NDDs.

The relevance of the PCB associated DNA methylation profiles described in our study to human disorders appears to reflect the evolutionary conservation of developmental events. This is evidenced by the top chromatin state enrichment: Bivalent TSS. Although all chromatin states are highly conserved between human and mouse, the bivalent TSS chromatin state, which represents 1.2% of the entire genome across all tissues and ~0.3% in a specific tissue, is substantially more evolutionarily conserved than the other 17 chromatin states (van der Velde et al. 2021). There are ~3000 bivalent genes in each fetal tissue, which are poised for either activation and repression, and many of them are lineage specific transcription factors that are repressed in the tissue assayed but expressed in others (Ngan et al. 2020; van der Velde et al. 2021). Given the tissue specific nature of the identified bivalent chromatin state, future research into the effects of PCB exposure on DNA methylation profiles in placenta and brain would benefit from examining specific regions and cell populations through sorting or single-cell sequencing.

Taken together, these findings demonstrate that a human relevant PCB mixture results in differentially methylated genes in placenta that are also present in the developing the brain, which represents disruptions to cellular signaling pathways of relevance to both tissues. This suggests that the placenta, a typically discarded birth by-product, contains DNA methylation profiles relevant to brain consequences of prenatal PCB exposure induced alterations in the brain and associated NDDs. Future research focused on DNA methylation profiling of human term placenta with measured PCB exposures, and maternal blood-derived cell-free fetal DNA released from placental trophoblast (Alberry et al. 2007), could potentially lead to the development of informative biomarkers and enable early diagnosis of prenatal exposures and early intervention of associated NDDs.

## Author’s Contributions

JML, PJL, RJS, BIL, KEN, and AEV designed the study. JML, PJL, and RJS acquired funding for the study. JML and PJL supervised the project. KEN, AEV, and DHY performed the mouse work. KEN and BIL performed the DNA and RNA isolations. BIL performed the bioinformatic analyses. BIL interpreted the results and wrote the manuscript with intellectual contributions from JML. JML, PJL, and KEN edited the manuscript. All authors reviewed and approved the final manuscript. We also thank Dr. Kimberly Keil Stietz and Dr. Annie Ciernia for input related to the mouse work.

---

**Supplementary Figure 1**: Heatmap of unique DMR gene symbol mappings that are shared between the female and male placenta-brain overlaps and the top enrichments from previously published RNA-seq datasets.

**Supplementary Table 1**: Significant (empirical *p* < 0.05) DMRs from sex stratified pairwise contrasts of prenatal PCB exposed placenta and fetal brain.

**Supplementary Table 2**: Significant (*p* < 0.05) slimmed GO terms for the prenatal PCB exposure DMRs.

**Supplementary Table 3**: Significant (*p* ≤ 0.01) transcription factor motif enrichments for the prenatal PCB exposure DMRs.

**Supplementary Table 4**: Enrichment testing results for the prenatal PCB exposure DMRs within CpG and gene region annotations.

**Supplementary Table 5**: Enrichment testing results for the prenatal PCB exposure DMRs within forebrain tissue from an 18 chromatin state model of mouse embryonic development.

**Supplementary Table 6**: Annotations for the 210 gene symbol mappings that overlap between all pairwise contrasts of prenatal PCB exposure DMRs.

**Supplementary Table 7**: Significant (*q* < 0.05) slimmed GO terms, Panther pathways, and GEO RNA-seq disease and drug signature enrichments from a meta p-value analysis of genes symbol mappings from the sex stratified overlaps of placenta and brain.

