## Supplementary figures and images for "Genome-Wide DNA Methylation Profiles of Neurodevelopmental Disorder Genes in Mouse Placenta and Fetal Brain Following Prenatal Exposure to Polychlorinated Biphenyls"

### Supplementary Figure 1

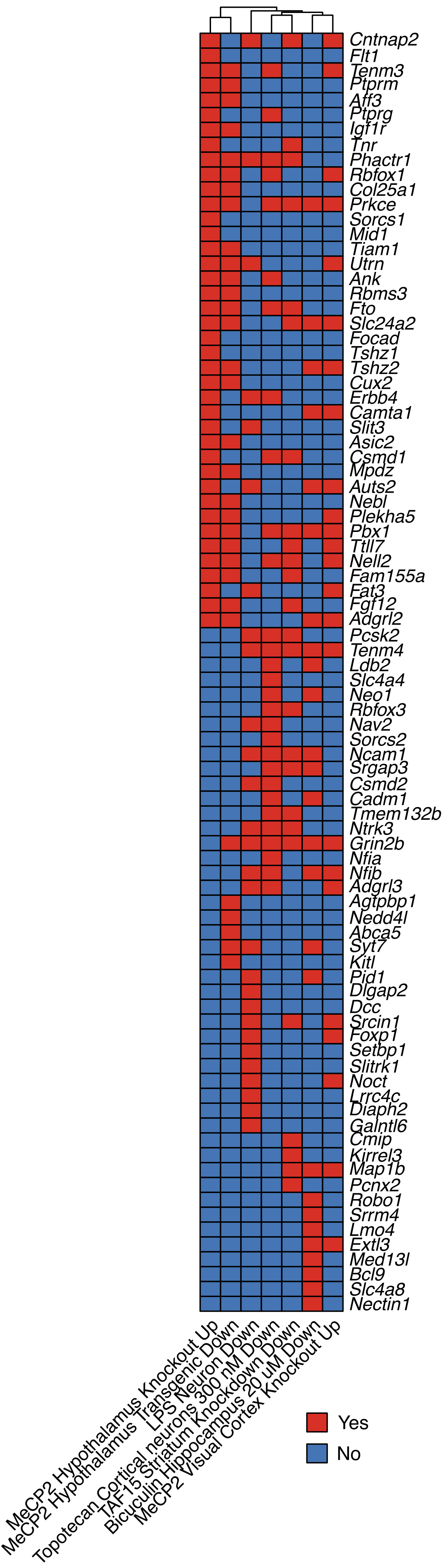
